# Deep-Plant: a supervised foundation model for plant regulatory genomics

**DOI:** 10.64898/2026.04.06.716755

**Authors:** Ahmed Daoud, Soumyadip Roy, Haoxuan Zeng, Xinyu Bao, Zhenhao Zhang, Jiakang Wang, Paul Parodi, Anireddy SN Reddy, Jie Liu, Asa Ben-Hur

## Abstract

Large-scale sequence-to-function deep learning models have demonstrated unparalleled ability to model biological sequences and have revolutionized the field of regulatory genomics. However, the majority of such efforts have centered on human and mammalian systems, leaving plant regulatory genomics comparatively underexplored. To address this gap, we introduce Deep-Plant, a supervised foundation model trained to predict chromatin state directly from genomic sequence. In contrast to large language models, which are trained in a selfsupervised manner using sequence alone, our model is trained to predict chromatin state across tissues and conditions. Training the model on a large collection of genome-wide experiments including DNA accessibility, transcription factor binding, and histone modifications, provides it with added biological context beyond the sequence itself. We demonstrate that the resulting model is an effective platform for developing accurate models of regulatory activity relevant to gene expression and active enhancers, exhibiting large improvements in speed, accuracy, and interpretability over the complementary approach of fine-tuning DNA language models. Deep-Plant models are available in Arabidopsis and rice, and work well as a building block for sequence modeling in related species such as corn. Together, these results establish supervised, chromatin-informed foundation models as a practical and effective paradigm for regulatory sequence modeling in plants.

## 1 Introduction

Deciphering how DNA sequence gives rise to gene regulatory activity in complex eukaryotic genomes, ranging from transcription factor binding and chromatin accessibility to histone modifications and gene expression, remains a central challenge in genomics. This challenge, often referred to as the *sequence-to-function* problem [1], aims to leverage modern deep learning in conjunction with large datasets to uncover how *cis*-regulatory elements control developmental programs, stress responses, and species-specific traits. Understanding the *cis*-regulatory code is critical both for basic biology and for applications in biotechnology and agriculture [2, 3].

In recent years, deep learning models trained on thousands of genome-wide datasets have demonstrated an ability to model the complexities of gene regulation in mammalian systems at nucleotide resolution, providing a detailed view of diverse genomics and epigenetics data, including transcription factor binding, DNA accessibility, and gene expression [4, 5]. Such models are complementary to DNA language models, and in practice, provide a better starting point for modeling tasks in regulatory genomics [6, 7].

The paradigm of foundation models, large-scale models typically pre-trained in a self-supervised fashion on massive sequence datasets and followed by fine-tuning to downstream tasks, has emerged as a powerful approach in regulatory genomics. Transformer-based DNA language models have been trained across hundreds of species and have demonstrated strong performance in sequence-based representation learning and transfer learning for genomic tasks [8, 9, 10, 11, 12, 13]. DNA language models have also been developed for plants, demonstrating the benefit of training models that specifically represent the characteristics of plant genomes [14, 15].

However, regulatory function is not encoded in DNA sequence alone. In eukaryotic genomes, DNA is embedded in chromatin, and gene regulation is mediated through chromatin accessibility, transcription factor binding, and histone modifications that vary across tissues and conditions [16, 17]. Models trained on sequence alone must therefore infer regulatory activity without explicit information about these regulatory layers. In contrast, modeling approaches that explicitly incorporate chromatin state provide direct supervision related to regulatory function, offering the potential for more accurate, interpretable, and biologically grounded representations of the *cis*-regulatory code.

Sequence-based models that are informed by large-scale chromatin state or expression datasets provide an effective alternative to DNA language models by directly modeling regulatory activity [18, 5, 4, 19]. By integrating these additional layers of information, such models provide an effective platform for downstream modeling problems, much the same way that DNA LLMs, but often with improved accuracy, enhanced interpretability and faster training [7, 6]. While such models are widely available in mammalian genomes, they are largely missing in plant systems [20, 3, 21]. Although deep learning models are available in plants to model specific aspects of chromatin state such as DNA accessibility or TF binding [22, 23, 24], these models do not match the scale or breadth of the models available for mammalian genomes. With the recent availability of ENCODE-style resources of uniformly processed plant epigenomic data, such as ChIP-Hub [25], assembling the datasets required to train large-scale chromatin-informed models has become feasible. And indeed, the lab behind ChIP-Hub has recently analyzed histone modification datasets using deep learning [26], an effort that is complementary to the work presented here.

Taken together, these observations point to a clear gap in current approaches in plant regulatory genomics. To address this gap, we introduce Deep-Plant, a supervised foundation model designed specifically for plant genomes that directly models chromatin state from DNA sequence. The model is trained to predict DNA accessibility, transcription factor binding, and histone modifications using large datasets composed of 2,8535 Arabidopsis samples and 350 rice samples retrieved from ChIP-Hub. We demonstrate that Deep-Plant achieves improved accuracy and computational efficiency compared to DNA language models on downstream tasks, including the prediction of gene expression and active enhancers directly from sequence. Furthermore, our results show that the representations learned by Deep-Plant transfer across dicots and monocots, facilitating the development of deep learning models across diverse plant species. This establishes Deep-Plant as a valuable resource for interpreting plant genomes at a nucleotide level that is complementary to DNA language models.

Overall, this work brings the supervised foundation-model paradigm to plant regulatory genomics and establishes chromatin-informed modeling as a scalable and biologically grounded approach for interpreting plant genomes.

## 2 Results

We developed Deep-Plant, a sequence-based supervised foundation model for plant regulatory genomics, trained on extensive chromatin state datasets from Arabidopsis and rice (Figure 1). Leveraging data from ChIP-Hub [25], the model incorporates genome-wide chromatin state information from 2,835 experiments in Arabidopsis and 350 in rice, providing information on DNA accessibility, histone modifications, DNA binding, and DNA methylation. The architecture of Deep-Plant is composed of three key components: a convolutional backbone, a transformer encoder block, and an attention-based pooling mechanism. This design enables the model to capture both local sequence motifs and long-range dependencies critical to regulatory interactions. The convolutional module extracts motif-level and mid-range features through stacked residual and pooling layers. These features are then processed by a multi-layer transformer encoder, which models long-distance regulatory interactions across the sequence. An attention pooling layer then summarizes the embeddings into a representation, which is passed through a multi-layer prediction head producing predictions of genome-wide epigenomic signals. Using a relatively short 2.5kb DNA sequence as input, the model is optimized for the compact genomes of Arabidopsis and rice, which are characterized by higher gene density and shorter intergenic regions compared to mammalian systems. Deep-Plant is first pre-trained to predict chromatin state, including DNA accessibility, histone modifications, DNA-binding events, and DNA methylation. The model is then fine-tuned for downstream tasks related to gene regulation including prediction of gene expression and enhancer activity, to showcase its utility for plant regulatory genomics.

**Figure 1:**
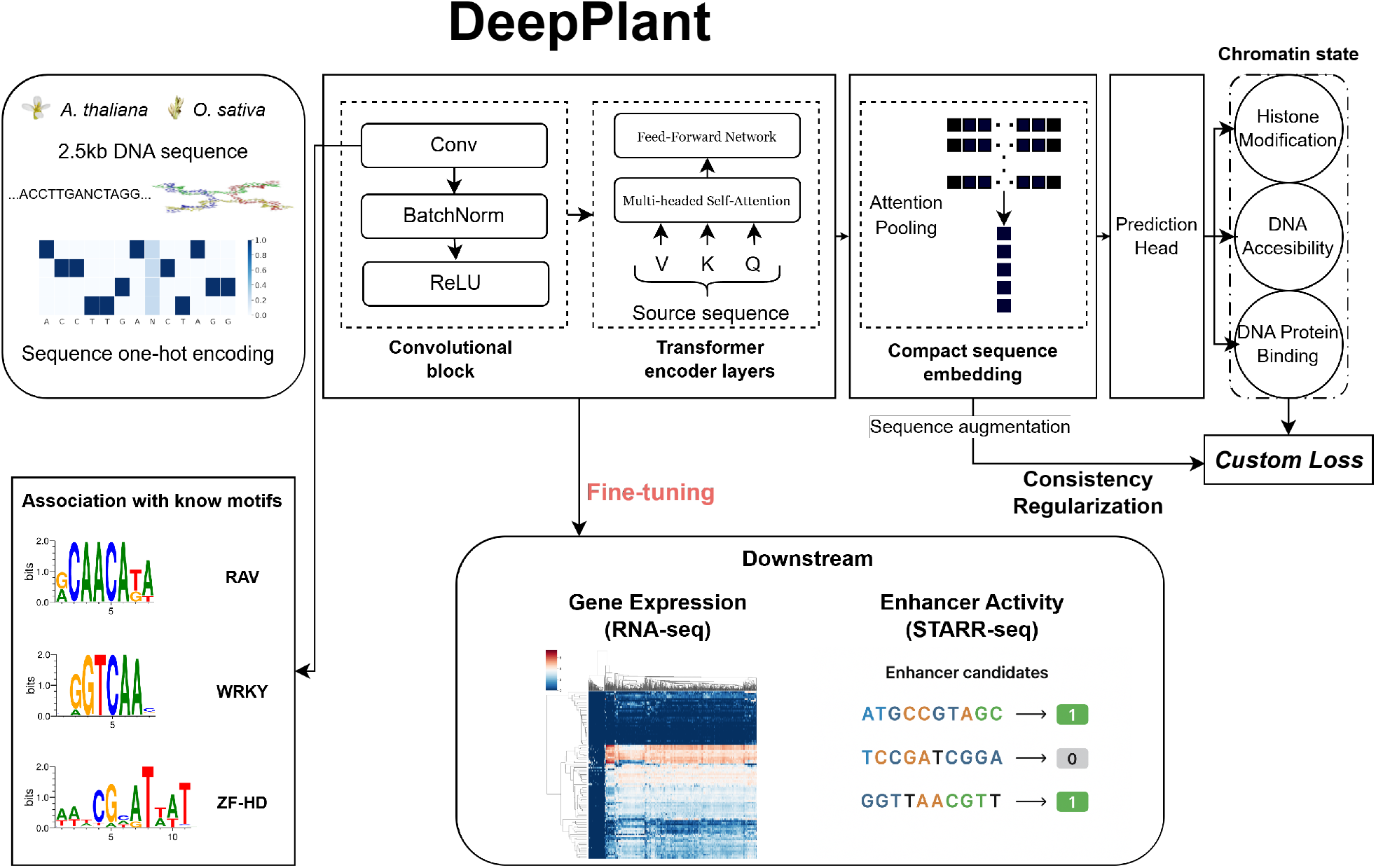
Schematic overview of Deep-Plant. The model processes DNA sequences of length 2.5 kb from Arabidopsis and rice by an architecture that integrates convolutional layers, transformer encoder layers. The prediction head outputs chromatin state features, including histone modifications, DNA accessibility, and transcription factor binding. Fine-tuning the model enables downstream applications such as gene expression and enhancer activity prediction.

### Chromatin state prediction

To evaluate the ability of Deep-Plant to model chromatin state, we trained the model using 2.5kb sequence windows to predict normalized read coverage within the central 200bp of each window. This ensures that the model focuses on the most biologically relevant region while leveraging flanking sequence context to capture regulatory interactions. Training was performed using a custom loss function based on the Poisson loss, which is well-suited for modeling count-based genomic data. The training data, derived from ChIP-Hub [25], included a comprehensive collection of genome-wide chromatin state datasets from Arabidopsis and rice. These datasets include DNA accessibility (DNase I-seq and ATAC-seq), transcription factor binding (TF ChIP-seq), and histone modifications (ChIP-seq). A detailed summary of the datasets is provided in Table 1 and Supplementary Table 1.

**Table 1:** Epigenomic data in Arabidopsis and rice: We collected all available data for Arabidopsis and rice in ChIP-Hub and averaged replicates. For DNA-binding proteins we list the overall number of datasets along with the number of unique proteins represented.

| Experiment Type | Arabidopsis | rice |
| --- | --- | --- |
| DNA-binding proteins | 1,547 (762 unique) | 74 (28 unique) |
| Histone and Histone-related | 1,119 | 271 |
| DNase and ATAC-seq | 164 | 5 |
| DNA-methylation | 5 | 0 |

We first evaluated the performance of Deep-Plant in predicting chromatin states across species and regulatory contexts. In Arabidopsis, the model achieved an average Pearson correlation of 0.680 between predicted and experimental read coverage across all chromatin state datasets, while in rice, it attained a mean Pearson correlation of 0.688 (Figures 2A, B). Notably, Deep-Plant demonstrated particularly high accuracy in experiments related to histone modifications and DNA-binding proteins, underscoring its ability to capture complex regulatory interactions encoded in the sequence. Additionally, our analysis revealed a positive correlation (*R* = 0.516) between Deep-Plant’s prediction performance and the number of peaks in the experimental BED files. The model exhibited superior accuracy on datasets with high signal density, whereas performance declined for experiments with sparse level of signal (Supplementary Figure 1).

**Figure 2:**
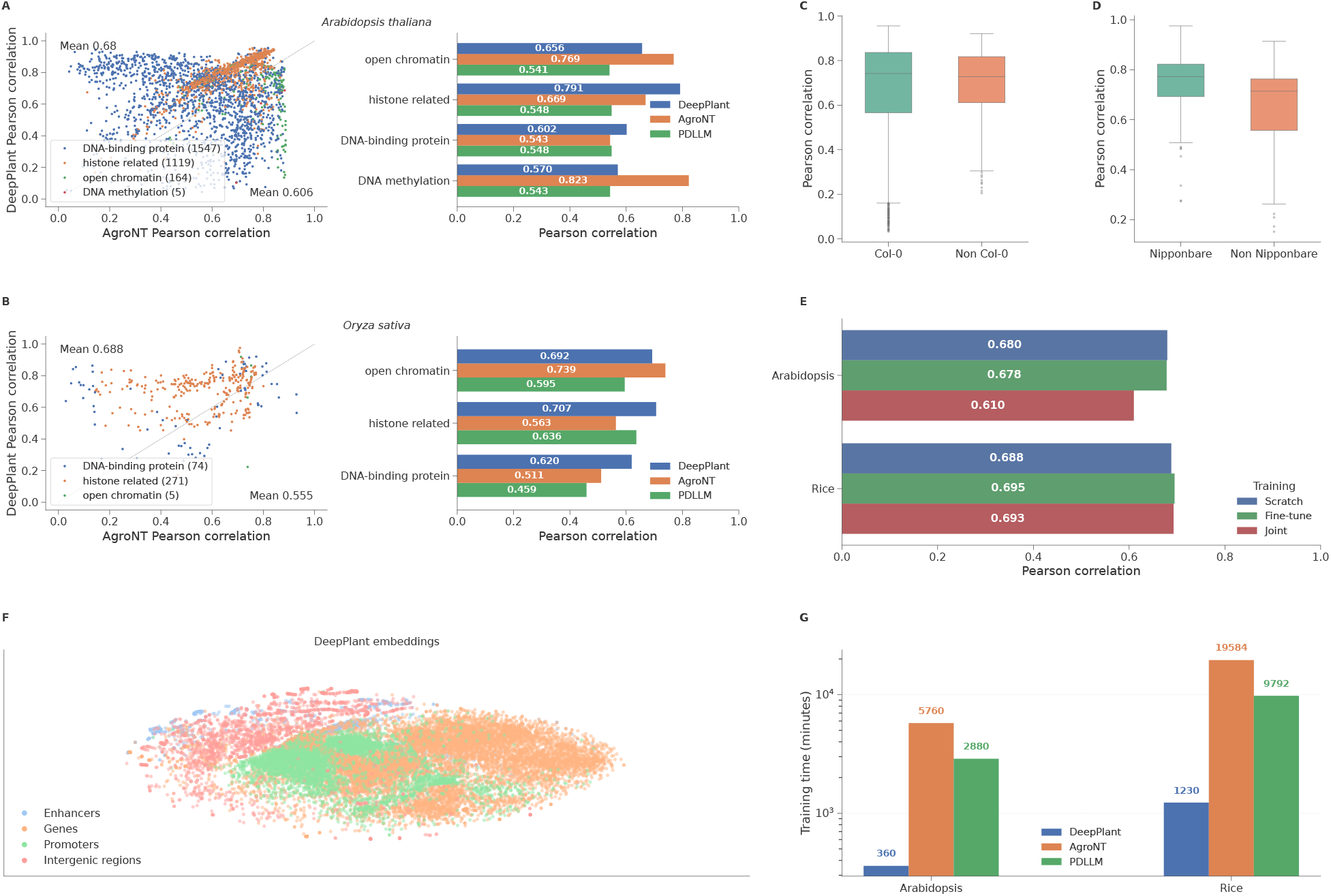
Comparative performance of Deep-Plant across plant species in predicting chromatin state. (A) Model performance on Arabidopsis across chromatin-related experiments (numbers in parentheses denote the number of experiments in each group). (B) Performance in rice across chromatin-related experiments. (C) Intraspecies generalization in Arabidopsis. Boxplot comparison of Pearson correlation values for Arabidopsis experiments grouped by accession (Columbia-0 vs. non–Columbia-0). (D) Intraspecies generalization in rice. Boxplot comparison of Pearson correlation values for *Oryza sativa* experiments grouped by accession (Nipponbare vs. non–Nipponbare) (E) Comparison of training time per epoch across foundation models. Bars show the average time (in minutes) required to complete one training epoch for Deep-Plant, AgroNT, and PDLLM in Arabidopsis and rice. (F) Evaluation of cross-species training strategies. Barplots comparing the average Pearson correlations in Arabidopsis (top) and Rice (bottom) across three training approaches: training from scratch, cross-species fine-tuning, and joint training. G) Deep-Plant embeddings reveal functional organization of regulatory DNA sequences in Arabidopsis. t-SNE visualization of 2,500 bp sequence embeddings generated by Deep-Plant.

To contextualize these results, we compared Deep-Plant performance to two recently developed plant DNA language models, AgroNT [14] and PDLLM [15]. Both serve as strong baselines as general-purpose pretrained models designed for downstream genomics applications. Similarly to Deep-Plant, these models rely solely on DNA sequence as input but differ in architecture and training strategies. Deep-Plant consistently outperformed both baselines (Figure 2A,B). We observe that AgroNT performed better on chromatin accessibility tasks, likely due to its ability to model detailed sequence context. However, Deep-Plant performed better overall, and its performance gains were particularly pronounced for experiments relating to histone modifications and DNA-binding proteins. We hypothesize that the improved performance of Deep-Plant in this context is due to its ability to model the relationships between motifs detected by its convolutional layers. In the Supplementary Information, we show that the first convolutional layer in Deep-Plant learns biologically relevant features matching known transcription factor motifs in Arabidopsis (Supplementary Figure 2).

#### Generalization to non-reference accessions

Next we assessed Deep-Plant prediction performance across accessions of Arabidopsis and rice to evaluate its ability to generalize to nonreference genomes. In Arabidopsis, we compared the model’s performance on the reference Columbia-0 (Col-0) accession to non-Col-0 accessions (Figure 2C). The dataset comprised 2,535 Col-0 experiments and 290 experiments from non-Col-0 lines. Deep-Plant produced similar correlation scores across both groups, achieving an average Pearson correlation of 0.678 for Col-0 and 0.693 for non-Col-0 lines, demonstrating robust generalization across Arabidopsis accessions.

In rice, we compared prediction performance on the reference genome of *Oryza sativa* Japonica Group (Nipponbare) to other rice accessions (Figure 2D). This dataset included 117 Nipponbare experiments and 233 experiments from non-Nipponbare accessions. Unlike Arabidopsis, Deep-Plant exhibited higher performance on the Nipponbare genome (Pearson correlation = 0.740) compared to non-reference accessions (Pearson correlation = 0.661).

We hypothesize that the disparity in generalization across accessions reflects differences in evolutionary history and genome architecture. Arabidopsis accessions exhibit relatively conserved genome structure, with limited structural variation [27]. In contrast, rice cultivars are characterized by high levels of structural variation [28], shaped by extensive domestication, adaptation, and breeding. For example, Schatz et al. found that approximately 9% of the IR64 indica genome assembly fails to align to the Nipponbare reference genome, highlighting the substantial structural divergence between rice subspecies [29]. While deep learning models like Deep-Plant are generally robust to single nucleotide polymorphisms (SNPs), they are less equipped to account for large-scale structural variations, which likely contribute to the observed differences in prediction performance across rice accessions.

#### Generalization across species

To investigate the transferability of genomic features between plant species, we evaluated three distinct training strategies: training from scratch (single-species), cross-species fine-tuning (transfer learning), and joint training with fully shared parameters (Figure 2E). Following Basenji2 [30], we used a shared architecture with species-specific prediction heads, a strategy previously shown to improve human-mouse joint training. For *Arabidopsis thaliana*, the model trained from scratch established a strong baseline with a median Pearson correlation of 0.680. Fine-tuning a model pre-trained on rice (OS *→* AT) yielded nearly identical performance (0.678). However, joint training with fully shared filters resulted in a significant performance drop to 0.610. In contrast, results for rice remained robust across all strategies, with fine-tuning, shared joint training, and non-shared joint training achieving correlations of 0.695, 0.693, and 0.685 respectively, comparable to the scratch baseline of 0.688. The decline in Arabidopsis performance in joint training with shared filters suggests that the use of shared filters in the joint architecture constrains the learning of Arabidopsis-specific patterns, likely pointing towards a divergence in epigenomic regulatory motifs between Arabidopsis and rice that cannot be optimally captured by a single shared encoder.

#### Deep-Plant is much faster than DNA LLMs

Not only does Deep-Plant achieve strong predictive accuracy, but it is also faster to train than fine-tuning DNA language models. To assess computational efficiency, we compared the average training time per epoch for Deep-Plant to fine-tuning AgroNT and PDLLM on a similar hardware setup. As shown in Figure 2G, Deep-Plant requires substantially less time while maintaining comparable or superior predictive accuracy. Owing to its hybrid architecture that combines convolutional feature extraction with transformer-based context modeling, Deep-Plant achieves an effective balance between computational efficiency and representational power, offering a practical advantage for large-scale chromatin-state prediction tasks.

#### The Deep-Plant embedding space

We pre-trained Deep-Plant using a consistency-regularization framework that enforces the production of stable embeddings across perturbed views of the same input sequence. By operating at the representation level, this objective enhances the model’s robustness to biologically plausible sequence augmentations, differing from approaches such as Basenji2 [30], which impose consistency directly on predictions. Consequently, Deep-Plant learns a representation space in which sequences with similar regulatory functions are embedded in close proximity. To examine how Deep-Plant represents different types of regulatory sequences in its embedding space, we extracted representations for 2,500 bp genomic segments from Arabidopsis enhancers, promoters, genes, and intergenic regions. The embeddings generated by Deep-Plant were projected into two dimensions using t-SNE, and the resulting scatterplot revealed a spatial organization that reflects the biological relationships among sequence classes (Figure 2F). Promoters and genes clustered tightly at the core of the embedding space, forming a dense central region. Intergenic sequences surrounded this core, delineating its boundaries and occupying the outer areas of the plot. Enhancers appear either adjacent to intergenic regions or along the periphery of the gene–promoter cluster, indicating that Deep-Plant recognizes them as functionally intermediate between genic and non-genic regions. This clear clustering pattern suggests that Deep-Plant captures intrinsic regulatory similarities and differences directly from DNA sequence context, even without explicit positional or functional supervision.

### 2.1 Using Deep-Plant to predict gene expression

To evaluate the effectiveness of Deep-Plant on downstream regulatory tasks, we fine-tuned the pretrained models to predict log transformed gene expression levels in *Arabidopsis thaliana* and *Oryza sativa*. Model performance was assessed using the Pearson and Spearman correlation coefficients and benchmarked against AgroNT and PDLLM (Figure 3A). Deep-Plant achieved strong predictive accuracy in both species. In Arabidopsis, it reached Pearson and Spearman correla-tions of 0.748 and 0.742, outperforming AgroNT (0.465 / 0.465) and PDLLM (0.508 / 0.513). In rice, Deep-Plant attained correlations of 0.781 (Pearson) and 0.737 (Spearman), again surpass-ing AgroNT (0.369 / 0.336) and slightly outperforming PDLLM (0.690 / 0.710).

**Figure 3:**
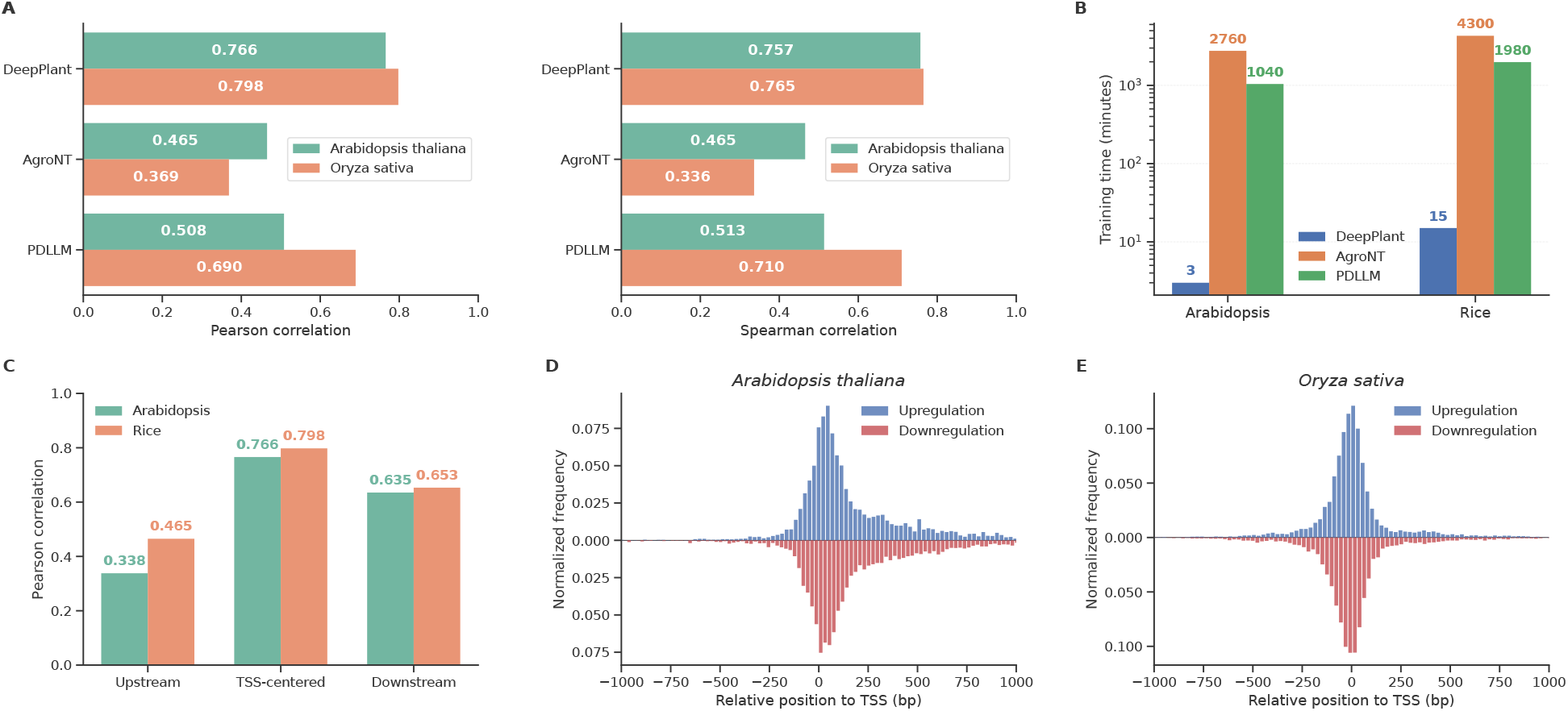
Gene expression prediction with fine-tuned Deep-Plant. (A) Pearson and Spearman correlations between predicted and observed gene expression averaged across 3,142 experiments; and results for rice averaged across 450 experiments. (B) Per-epoch training time comparison between Deep-Plant, AgroNT, and PDLLM. (C) Performance comparison across three 2.5 kb windows relative to the TSS (upstream, TSS-centered, downstream). (D) Normalized frequency of ISM-identified mutations relative to the TSS in Arabidopsis, separated into mutations causing upregulation (top) and downregulation (bottom). (E) Normalized frequency of ISM-identified mutations relative to the TSS in rice, separated into mutations causing upregulation (top) and downregulation (bottom).

Consistent with our observations on the chromatin-state task, fine-tuning Deep-Plant for gene expression prediction was 100x times faster than fine-tuning the large language models (Figure 3B).

#### Gene regulation in Arabidopsis and rice is TSS-proximal

Unlike mammals, the compact genomes of Arabidopsis and rice feature a highly condensed, TSS-proximal regulatory landscape [31, 32, 33, 34]. To evaluate the contribution of the regions upstream and downstream of the TSS, we fine-tuned Deep-Plant on windows 2.5 kb upstream of the TSS, 2.5 kb centered around the TSS, and 2.5 kb downstream windows (Figure 3C). In both species, TSS-centered sequences yielded the highest predictive performance, followed by downstream and then upstream regions, indicating that key regulatory signals are concentrated near and just downstream of the TSS [14]. We corroborated this at the nucleotide level through *in silico* mutagenesis (ISM) across the TSS-centered sequences. In Arabidopsis, influential mutations are strongly enriched around the TSS and decay more slowly on the downstream side (Figure 3D), similar to observations in previous work [32, 14].A similar, though less downstream-skewed, pattern emerged in rice (Figure 3E). This dense regulatory neighborhood extends downstream into the 5’ UTR and first intron, serving as essential hubs for transcriptional initiation and intron-mediated enhancement, especially in Arabidopsis [35, 36]. Together, our results show that TSS-proximal regions are the primary drivers of plant transcriptional output.

#### A case study: the DREB1 gene family in Arabidopsis

To investigate regulatory mechanisms, we performed an *in silico* mutagenesis (ISM) on the cold-inducible DREB1 gene cluster (AT4G25470-AT4G25480-AT4G25490) in *Arabidopsis thaliana* [37]. While extensive research has been performed on the regualtion of the DREB1 gene family to identify their regulatory elements, this research focused on regulators in the promoter regions of these genes [38, 39, 40]. Our analysis reveals that strong regulatory signals are also prevalent in the 5’ UTRs of these genes, where they overlap with open chromatin and ChIP-seq-supported binding sites. We visualized the ISM scores for the three genes at different cold-stress timepoints (Figure 4) and highlighted the mutations with the strongest impact on predicted expression and transcription factor binding sites (see Methods section 4.3 for details). Notably, six of the eight most impactful regulatory motifs were within the 5’ UTR, underscoring its central role in governing gene expression in compact genomes. The ISM results shown in Figure 4 show dynamics in the response to cold, particularly in the role of the evening element motif. Furthermore, the TFs identified in our analysis are in agreement with TFs that are known to regulate the DREB1 genes [38, 39, 40]. While the precise mechanisms governing DREB1 expression through UTR elements remain unresolved, Deep-Plant provides a high-resolution framework for prioritizing candidate regulatory elements. This enables targeted experimental validation and can accelerate the dissection of transcriptional control in plant stress response.

**Figure 4:**
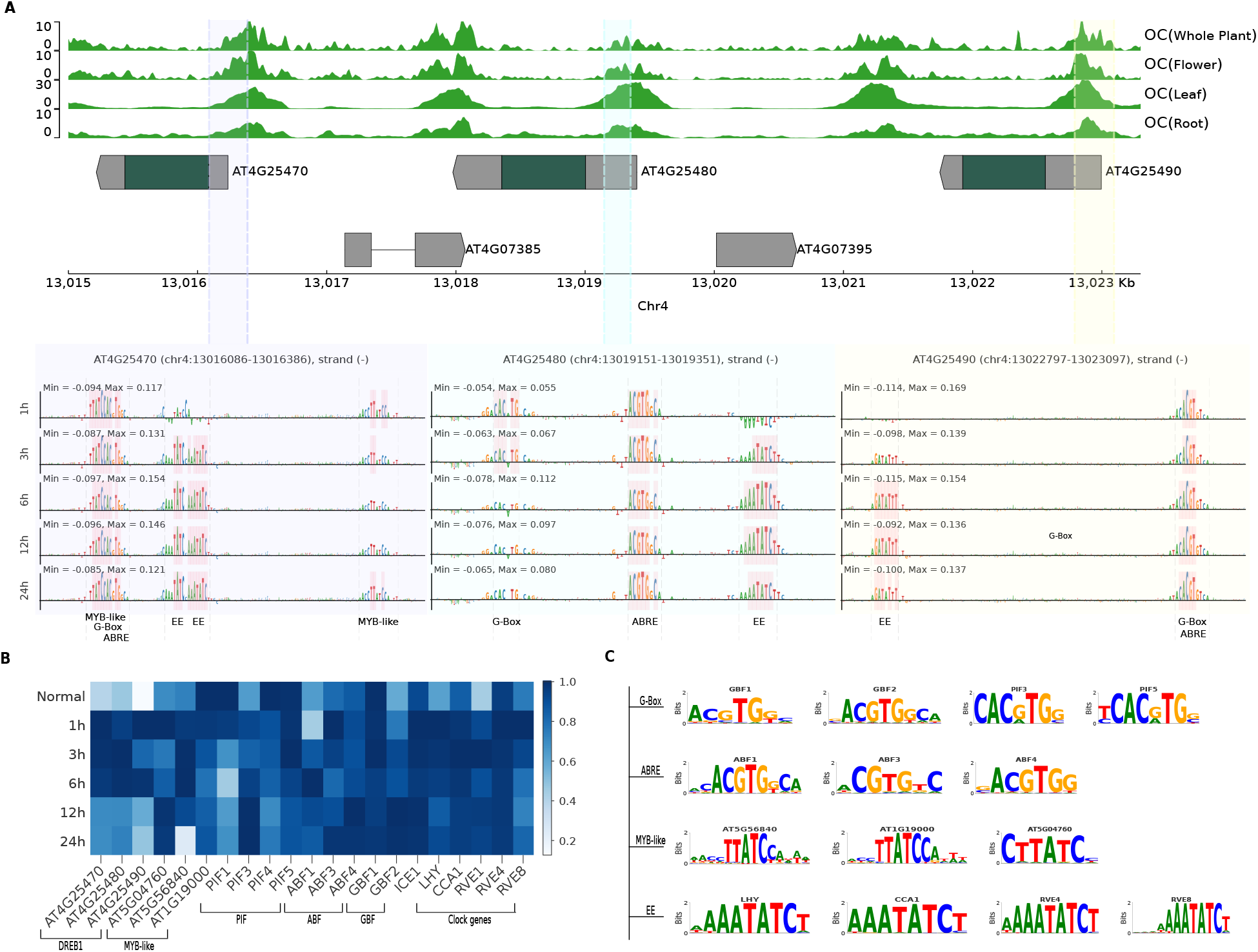
Regulatory Landscape and In-Silico Mutagenesis (ISM) Analysis of the Arabidopsis thaliana DREB1 Gene Cluster. (A) Genomic track view of the DREB1 cluster (AT4G25470, AT4G25480, AT4G25490) on chromosome 4. The top panels show open chromatin (OC) signals across different tissues (Whole Plant, flower, leaf, and root). Below, ISM identifies highly impactful regulatory motifs. Highlighted red bars indicate base pairs where mutations resulted in the highest absolute ISM scores, representing putative transcription factor (TF) binding sites critical for regulation across a 24-hour cold stress time course . (B) Heatmap showing the expression profiles of the DREB1 genes and their associated regulatory transcription factors under normal conditions and throughout a cold stress time course (1h, 3h, 6h, 12h, and 24h). Expression values are normalized for each gene. (C) Motif logos for TFs whose binding sites were identified as high-impact via ISM and disrupted by ISM-identified mutations.

### 2.2 Using Deep-Plant to predict enhancer activity

To further evaluate the capacity of chromatin-pretrained representations to capture functional regulatory activity, we fine-tuned the Arabidopsis Deep-Plant model to predict enhancer activity directly from sequence, using data derived from recent STARR-seq experiments [41], which provide direct measurements of enhancer activity. The fine-tuned Deep-Plant model achieved strong performance, with an area under the precision–recall curve (AuPRC) of 0.946, outperforming AgroNT (AuPRC = 0.881) and PDLLM (AuPRC = 0.832) (Figure 5A).

**Figure 5:**
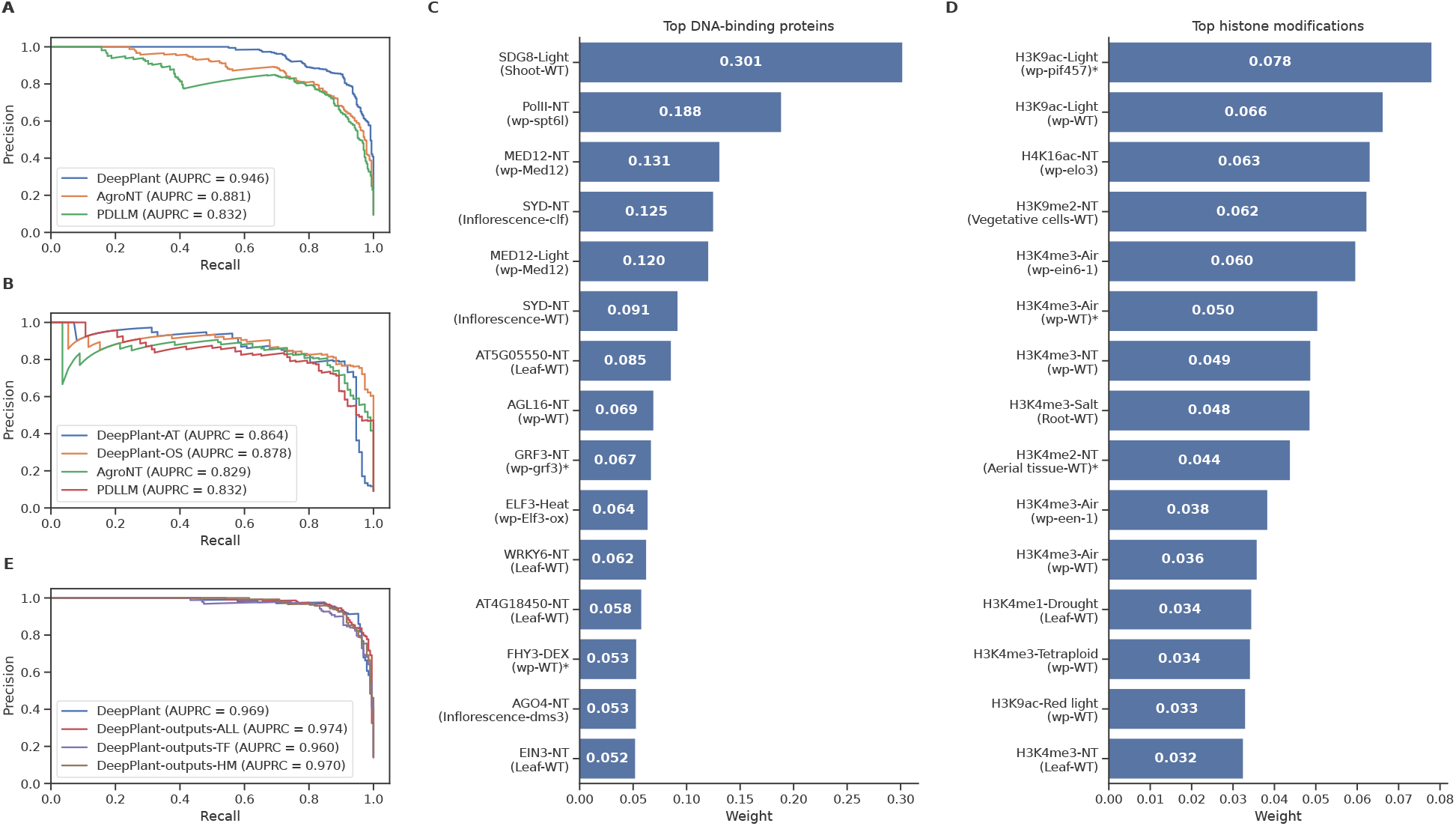
Enhancer activity prediction in Arabidopsis. (A) Precision–recall (PR) curves comparing Deep-Plant, AgroNT, and PDLLM in predicting Arabidopsis enhancer activity. (B) Precision–recall (PR) curves comparing Deep-Plant and Deep-Plant-outputs’ different flavor models. (C) Top ten weighted features for enhancer sequences derived from the logistic regression model using only DNA-binding features. (D) Top ten weighted features for enhancer sequences derived from the logistic regression model using only histone-related features. Treatments, tissues, and mutants are included to differentiate experiments; “NT” and “wt” stand respectively for no treatment and wild type. Experiments marked with * are non–Col-0 experiments. (E) Precision–recall (PR) curves comparing Deep-Plant-AT, Deep-Plant-OS, AgroNT, and PDLLM in predicting maize enhancer activity.

#### Deep-Plant identifies key epigenetic and transcriptional regulators associated with active Arabidopsis enhancers

To demonstrate the accuracy and interpretability of Deep-Plant’s chromatin state predictions, we trained logistic regression classifiers to identify high-confidence active enhancers (defined by DHS overlap and proximity to highly expressed genes; Methods 4.3). These classifiers use the chromatin predictions generated by Deep-Plant as input features. The resulting classifiers, collectively referred to as Deep-Plant-outputs, were constructed using different subsets of predicted targets (i) DNA-binding–related outputs, (ii) histone modification–related outputs, and (iii) the complete set of predicted epigenomic outputs.

Despite their simplicity, these models performed very competitively (AuPRC = 0.960-0.974) with a fine-tuned Deep-Plant model on the same dataset (AuPRC = 0.969) (Figure 5E), indicating that the chromatin state representation encodes information highly relevant to identifying enhancers. Furthermore, we examined the most informative epigenomic features contributing to enhancer classification. The analysis was conducted separately for DNA-binding and histone related features, allowing us to identify which regulatory signals were most informative for enhancer prediction in Arabidopsis.

Figure 5C highlights the DNA binding proteins driving model performance. The most heavily weighted factor, the histone methyltransferase SDG8, deposits H3K36me3 to antagonize Polycomb-dependent silencing [42]. The high predictive value of Pol II indicates these enhancers actively transcribe eRNAs [43]. Additionally, the model captures the structural mechanics of enhancer ac-tivity by prioritizing the nucleosome remodeler SYD [44] and the Mediator complex component MED12 [45, 46]. Finally, it identifies key developmental and stress regulators (GRF3, WRKY6, EIN3) [47, 48, 49] alongside known repressors (ELF3, AGL16, AGO4) involved in transcriptional and transposon silencing [50, 51, 52, 53].

Figure 5D highlights the histone modifications most predictive of enhancer activity. The model heavily weights acetylation marks like H3K9ac and H4K16ac, which drive chromatin accessibility and active regulation [54, 55]. It also prioritizes specific methylation marks, including the repres-sive mark H3K9me2 which represses some light-responsive genes [56], and various condition-specific H3K4me3 marks; while traditionally associated with promoters [57], their high predictive value here likely reflects the physical proximity of active enhancers to transcribed genes.

#### Deep-Plant identifies functional maize enhancers via cross-species transfer learning

To assess the cross-species generalization of Deep-Plant, we evaluated the capacity of our models’ representations to capture functional regulatory activity across divergent lineages. Specifically, we fine-tuned both the Arabidopsis and rice Deep-Plant models to predict enhancer activity in *Zea mays* directly from sequence, utilizing data derived from Oka et al. [58]. The rice-trained model achieved the highest overall performance (AuPRC = 0.881), surpassing the Arabidopsis model (AuPRC = 0.864) and outperforming AgroNT (AuPRC = 0.829) and PDLLM (AuPRC=0.832) (Figure 5E). The superior performance of the rice-pretrained model likely reflects the greater phylogenetic proximity between rice and maize, enabling better transfer of conserved monocot regulatory features despite the more comprehensive functional annotations of the Arabidopsis dataset.

## 3 Discussion

The development of Deep-Plant addresses a long-standing gap between the computational tools available for mammalian systems and those tailored for plants. Unlike approaches that rely primarily on unsupervised DNA language modeling, Deep-Plant builds on the supervised foundation model paradigm, directly integrating chromatin state information into sequence-based modeling. By leveraging a large-scale, multi-modal dataset encompassing DNA accessibility, transcription factor binding, and histone modification profiles, Deep-Plant learns sequence representations that enable more accurate and interpretable predictions of gene regulation than state-of-the-art DNA language models. This work demonstrates that supervised foundation models informed by chromatin context can outperform fine-tuned DNA language models in accuracy, speed, and biological interpretability.

One of the most impactful outcomes of this study is the demonstration that state-of-the-art genomic modeling does not require massive computational resources. By leveraging a hybrid architecture that combines convolutional layers for capturing local sequence motifs with transformer encoders for modeling long-range dependencies, Deep-Plant effectively balances local and global sequence information. This design enables the model to achieve superior performance while remaining significantly more computationally efficient than large DNA language models such as AgroNT and PDLLM. Notably, fine-tuning Deep-Plant is 10–100 times faster than finetuning these models, making it feasible to use commodity hardware. Furthermore, the inclusion of consistency regularization during pre-training played an important role in improving computational efficiency. By encouraging the model to produce stable predictions under input perturbations, this approach eliminated the need for computationally expensive sequence augmentation during fine-tuning, further enhancing the practicality of Deep-Plant for genomic applications.

Unlike black box deep learning models, Deep-Plant provides transparent insights into the cis-regulatory code of plants. The model’s convolutional filters autonomously learned biologically relevant features. Analysis showed that 98.83% of the filters matched known TF binding motifs.

The ability of Deep-Plant to generalize across Arabidopsis accessions highlights its potential to uncover conserved regulatory mechanisms in plant genomes, which are essential for understanding gene expression across diverse environmental contexts. The model’s performance on chromatin state datasets highlights its ability to accurately identify and characterize critical regulatory elements, including enhancers and transcription factor binding sites, which are integral to the control of gene expression. Notably, Deep-Plant also demonstrated strong cross-species generalization, as evidenced by its application to enhancer detection in corn, a crop with a significantly larger and more complex genome than Arabidopsis or rice. Furthermore, a model pre-trained on Arabidopsis and fine-tuned for rice chromatin state prediction achieved a Pearson correlation of 0.692 after only a single epoch of training. This result suggests that Deep-Plant captures fundamental regulatory principles shared across monocots and dicots, paving the way for applications in crops where experimental datasets are sparse or incomplete.

### Limitations and Future Directions

While Deep-Plant demonstrates excellent performance across a wide range of chromatin state prediction tasks, several limitations remain. First, the model’s reduced accuracy on rice accessions outside the Nipponbare reference highlights the need for training on more diverse datasets to enhance generalizability. Incorporating chromatin state data from indica and other subspecies could improve the model’s ability to capture the regulatory variation inherent to rice genomes. This disparity in performance reflects the challenges posed by the higher structural variation ob-served in rice compared to Arabidopsis. Addressing these challenges will require computational strategies specifically designed to account for structural variation, which is currently difficult for deep learning models to capture.

Second, the current architecture is optimized for compact genomes like Arabidopsis and rice, but further work is needed to assess its scalability to larger and more complex genomes, such as those of maize or wheat. These genomes exhibit greater complexity, including extensive repetitive elements and larger intergenic regions, which may require adaptations to the model architecture to ensure accurate predictions while maintaining the efficiency of the model.

Finally, while the supervised foundation model paradigm offers improved accuracy and interpretability, its reliance on large-scale labeled datasets can limit its applicability in species where chromatin state data are sparse or unavailable. Future research could explore hybrid approaches that combine supervised and self-supervised learning to leverage unlabeled sequence data while still incorporating chromatin context. Such strategies could enable the development of foundation models for non model plant species.

## 4 Methods

### 4.1 Chromatin state data

#### Data assembly and pre-processing

We downloaded the genomes of *Arabidopsis thaliana* Columbia (cv. Col-0) and *Oryza sativa* japonica group (cv. Nipponbare) from Ensembl plants [59]. We downloaded all available chromatin state data for those species from ChIPHub [25]; data was downloaded in read per genomic content (RPGC) normalized Bigwig format. After averaging replicates the resulting Arabidopsis compendium included 2,835 chromatin tracks, spanning DNA-binding proteins (n = 1,547), histone-related (n = 1,119), open chromatin (n = 164), and DNA methylation (n = 5). The rice dataset comprised of 350 experiments across DNA-binding proteins (n = 74), histone-related (n = 271), and open chromatin (n = 5).

To enhance the quality of the data, we excluded blacklisted regions from both genome sequences to remove regions prone to artifacts and noise. For Arabidopsis, blacklisted regions were identified using the ENCODE Blacklist tool [60], a resource specifically developed to exclude problematic regions of the genome that generate unreliable signals in high-throughput sequencing experiments such as ChIP-seq, DNase-seq, and ATAC-seq. The blacklist file for Arabidopsis was obtained from the project’s GitHub repository. For rice, we used published blacklist regions generated using GreenScreen [61]. Additionally, repetitive regions in both genomes were identified using RepeatMasker [62] and masked to minimize bias in downstream analyses.

We then applied several filtration steps to select qualified experiments from ChIP-Hub:

- Peak counts: We excluded experiments with less than 100 peaks in their respective bed files.
- Read-depth: We excluded experiments with more than 99% of the values in the bigwig file equal to 0.
- Missing data: We excluded experiments with missing data.
- Replicates: For experiments with multiple replicates, we calculated the average coverage read values to represent those experiments.

#### Input Data and Label Generation

Each genome sequence and read file was divided into 10 kb chunks for further processing. The 10 kb chunks were divided into train, test, and validation sets as described below. Within each 10 kb sequence and its corresponding read coverage data, we applied a sliding window approach to generate input and label data. The input to the model are sequence windows of length 2,500 bp, and the labels for each sequence were obtained by computing the average normalized read coverage in the center 200 bp of each sequence window. For each 10 kb chunk we slide a 2.5 kb window moving it in increments of 50 bp in the train set; in the validation and test set we used a 200 bp increment to obtain non-overlapping labeled windows.

To safeguard the training process from outlier values, we applied a transformation to the BigWig signal tracks, following the method used in Bassenji2 [30]. Specifically, we soft-clipped high values using the function 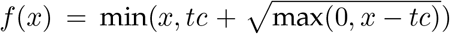 . This limits the contribution of values above a threshold *tc*, by incorporating only the square root of the residual *x* − *tc*. This approach aims to reduce the influence of extreme values unlikely to generalize across other genomic regions. We determined a consistent clipping threshold of *tc* = 50 for all experiments.

#### Train-Validation-Test split

We divided the genome into training, validation, and test sets in a way that ensures minimal sequence similarity across these sets. To group sequences into training, validation, and test sets and minimize sequence similarity across these sets, we ran BLAST [63] with epsilon set to 0.0001 for each 10kb chunk against its corresponding genome. Sequences that share sub-regions with sequence similarity of 70% and higher were grouped into distinct sets. The sub-region size was 400 bp for Arabidopsis and 1000 bp for rice. The 10kb chunks were then partitioned into training, validation, and test datasets and used to make the 2.5kb sequences.

The dataset contains 1,431,758 (sliding window of 50 bps) training, 45,125 (sliding window of 200 bps) validation, and 45,246 (sliding window of 200 bps) test sequences for the Arabidopsis genome and 4,466,694 (sliding window of 50 bps) training, 140,938 validation, and 140,589 test sequences for the rice genome.

### 4.2 Pre-training Deep-Plant

#### Model architecture

The Deep-Plant architecture includes three parts: (1) a block of convolutional and residual convolutional layers to extract the relevant sequence motifs from the DNA sequence using a genome-specific convolutional layers with a Rectified Linear Unit (ReLU) activation function; (2) six transformer encoder layers, and (3) an output head for chromatin state prediction (see Figure 6).

**Figure 6:**
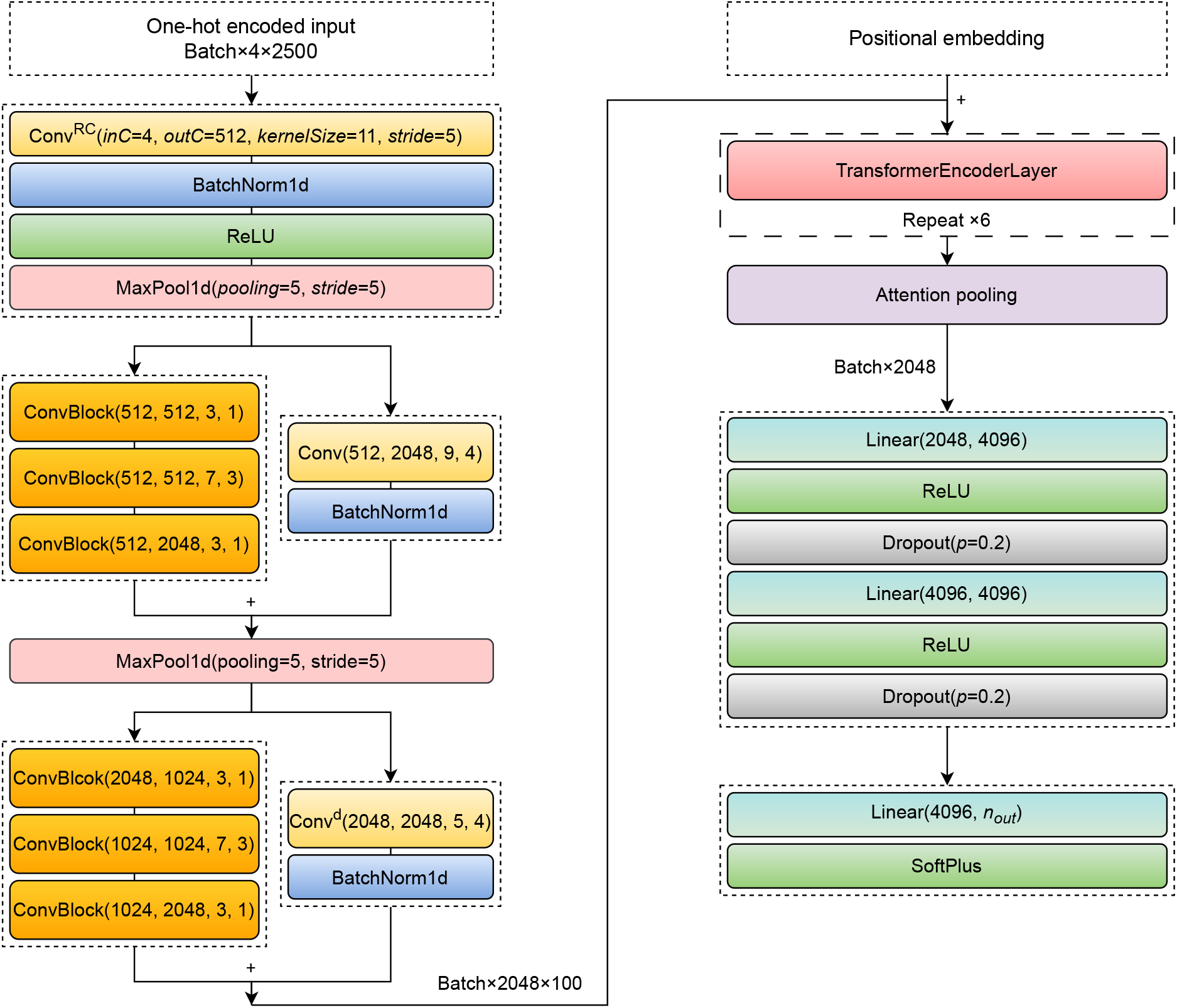
Deep-Plant model architecture. The network uses 1-dimensional convolutions parameterized by input channels (inC), number of filters (outC), kernel size (ks), and stride. **Conv**^*RC*^ denotes a reverse-complement convolutional layer where the output sums the convolutional products of both stranded (normal and reverse complement) filters. The **ConvBlock** consists of a convolution followed by a ReLU activation and Batch Normalization. **Conv**^*d*^ indicates a convolution with a dilation rate of 2. The architecture incorporates sinusoidal positional embeddings and a standard pytorch Transformer encoder layer (embedding dimension of 2048 with 8 attention heads). *p* represents the dropout probability. The output dimension, *n*_*out*_, corresponds to the number of chromatin state experiments: 2,835 for Arabidopsis and 350 for rice.

The first stage of the model is designed to extract relevant sequence motifs from the DNA input. First, a (genome-specific for joint training with non shared filters) convolutional head is applied, using 512 filters with a kernel (filter) size of 11, followed by a max-pooling layer (with a window size of 5 and a stride of 5). This captures local sequence patterns (motifs) and passes them to deeper layers. In this first convolutional layer, we employ reverse-complement parameter sharing, i.e., each convolutional filter is applied to the forward sequence and to its reverse-complement. This ensures that the model treats motifs and their reverse-complements equivalently, reflecting the biological fact that regulatory elements may occur on either DNA strand [64]. This increases strand-symmetry robustness and reduces redundancy by enforcing that complementary motif patterns map to the same learned representation, rather than learning separate filters for each orientation. Previous work on CNNs for DNA–protein binding prediction has shown that explicitly handling reverse-complement sequences (either via augmentation or via shared filters) improves performance and interpretability [65, 66]. Next, the model applies two convolutional blocks, separated by a max-pooling layer, each consisting of three standard convolutions and a skip connection convolution layer, similar to the ResNet-50 bottleneck block [67]. We used a dilated convolutional layer with skip connections in the second block. The final number of filters is increased by factors of 2 and 4 in these successive blocks. The convolutional block downsamples the input sequence length from 2500 bp to 100 bins, where each bin represents embeddings of a 25 bp segment of the DNA sequence.

The second stage of the architecture aims to capture long-range interactions. It uses attention-based encoders that are capable of modeling long-range relationships better than dilated convolution [18]. The encoder comprises six self-attention transformer blocks [68], each consisting of a multi-head self-attention layer and a feed-forward neural network layer. The self-attention mechanism of the *l*-th block operates as follows:

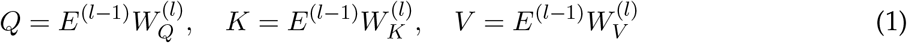

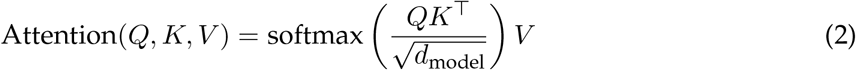

Here, *E*^(*l−*1)^ is the output of the (*l* −1)-th block, and *E*^(0)^ is the embedding produced by the convolutional blocks. *Q, K*, and *V* are the query, key, and value vectors respectively, each with dimension *N × d*_model_. The weight matrices 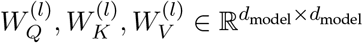, and the output projec-tion matrices 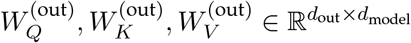.

Additionally, Deep-Plant splits the computation of self-attention into eight attention heads, allowing the model to jointly attend to information from different representation subspaces at different positions.

Next, an attention-pooling operation is then applied to obtain a single sequence representation vector *h*_*s*_ of dimension *d*_model_. This representation is passed to the final block, which consists of multiple non-linear fully connected layers and a genome-specific prediction head. A softplus activation function, *f* (*x*) = log(1 + *e*^*x*^), is applied to ensure positive outputs. The model predicts 2,835 epigenomic features for *Arabidopsis thaliana* or 350 for Oryza sativa.

#### Training and loss function

The model was trained to predict chromatin state profiles for two different plant species. Given a 2.5 kb DNA sequence, the model outputs the predicted average signal intensity for the central 200 bp region.

By operating at the representation level, this objective enhances the model’s robustness to biologically plausible sequence augmentations, differing from approaches such as Basenji2 [30], which impose consistency directly on predictions. Consequently, DeepPlant learns a representation space in which sequences with similar regulatory functions are embedded in close proximity.

The model was trained using the AdamW optimizer with a loss function that includes the Poisson and a custom consistency regularizer described below. We trained the model with a batch size of 16. Each sample was augmented six times, resulting in 96 distinct sequences per batch. To effectively increase the batch size and stabilize training, we accumulated gradients over four consecutive batches before performing a weight update. The learning rate was dynamically adjusted using a warmup–cosine decay schedule [69]. Specifically, the rate increased linearly from warmup _begin _lr = 1*e−* 4 to max_lr = 5*e−* 4 over warmup_steps = 5, and then decayed following a cosine annealing function to final_lr = 5*e−* 5. The scheduler parameters were empirically tuned based on the observed training and validation loss dynamics.

The training pipeline was optimized for distributed multi-GPU execution to accelerate convergence and improve throughput. To prevent overfitting and ensure stable generalization, we employed early stopping based on validation loss improvement over a predefined patience window. Additionally, a maximum epoch limit was enforced to manage computational resources efficiently. Ultimately, Deep-Plant was trained on two NVIDIA RTX 4090 (24 GB) GPUs for approximately two days.

To enforce invariance to biologically plausible transformations, we applied *consistency regularization*. The model was trained to minimize the pairwise cosine distances between the compact embeddings (*h*_*s*_) of an input sequence and its augmented variants (e.g., reverse complements and positional shifts). Unlike Basenji2 [30], which enforces invariance at the prediction level, our approach stabilizes representations directly at the embedding level. A key advantage of this strategy is that it improves downstream fine-tuning performance without the computational overhead of test-time sequence augmentation.

Formally, the overall training objective is defined as:

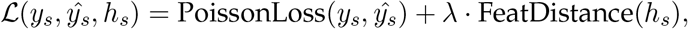

where *y*_*s*_ and ŷ_*s*_ denote the true and predicted chromatin signals, respectively, for the sequence *s*, and *h*_*s*_ = {*h*_1_, *h*_2_, …, *h*_6_} represents the sequence embeddings obtained from the attention pooling layer for the original and augmented sequences. The regularization-consistency term FeatDistance(*h*_*s*_) is computed as the mean pairwise cosine distances among all embeddings. The coefficient *λ* = 0.01 controls the strength of the regularization term and was selected empirically based on validation performance.

Note that we trained the model using the negative log likelihood with Poisson distribution of target loss or Huber loss, provided by PyTorch, and reported results using the Poisson loss since it performed slightly better than the Huber loss. We additionally evaluated the use of mean pairwise Euclidean distance, rather than cosine distance, for the regularization-consistency term. While this modification resulted in embeddings that appeared more visually separated in low-dimensional projections, it led to a slight decrease in overall model performance and increased variance across downstream tasks. Given this trade-off between visual interpretability and predictive stability, we retained cosine distance as the regularization metric.

#### Embedding visualization

To examine how Deep-Plant represents different types of regulatory sequences in its embedding space, we extracted representations for 2,500 bp genomic segments from Arabidopsis chromosomes 4 and 5. These included enhancers, promoters, genes, and intergenic regions. Focusing on chromosomes 4 and 5 ensured that most of the analyzed sequences originated from the model’s validation and test sets, minimizing overlap with training data. Intergenic sequences were defined as 2,500 bp segments located at least 5,000 bp away from any annotated enhancer, promoter, or gene.

Deep-Plant’s sequence embeddings were obtained from the attention pooling layer of the model. Each 2,500 bp sequence was transformed into a high-dimensional vector, which was subsequently reduced to two dimensions using t-distributed Stochastic Neighbor Embedding (t-SNE). The two-dimensional embeddings were visualized using a Seaborn scatterplot, where each point corresponded to a single sequence and was colored according to its class label.

### 4.3 Task-specific downstream data and models

#### Fine-tuning Deep-Plant for gene expression

To evaluate the transferability of the pretrained representations, we fine-tuned Deep-Plant to predict gene expression. Gene expression datasets for rice and Arabidopsis were obtained from the Plant Public RNA-seq Database (PPRD) [70]. For each species, samples with low-quality mapping (uniquely mapped read rate *<* 70%) were excluded. In addition, only rice samples belonging to the Japonica group and Arabidopsis samples of the Col-0 ecotype were retained for analysis. Expression matrices were normalized using FPKM values. To reduce technical variability, we identified biological replicates based on shared metadata attributes, including project, sample title, tissue, genotype, and treatment. Replicate samples were assessed for concordance using Pearson correlation. When pairwise correlations among replicates exceeded 0.95, they were merged by averaging their expression profiles. A second merging step further consolidated samples with highly similar titles, differing only by numeric suffixes, under the same correlation threshold. Following quality control and replicate merging, gene-level annotations were integrated using reference genome annotations for each species. Only genes with valid genomic coordinates and strand information were retained. The final expression matrices, aligned to these curated gene sets, were used for downstream analyses.

Genes were divided into training, validation and test sets according to their overlap with the corresponding Deep-Plant training, validation, and test sets. For Arabidopsis, the dataset comprised of 24,450 training, 3,908 validation, and 3,843 test genes, each associated with expression measurements across 3,142 experimental conditions. For rice, the dataset included 44,433 training, 5,764 validation, and 5,602 test genes, with expression values spanning 450 conditions or tissues. Fine-tuning was conducted independently for each species using the pretrained model as initialization. Model performance was evaluated on the held-out test set by computing both Pearson and Spearman correlation coefficients between the predicted and observed expression values across all experimental conditions and reporting the average correlation across genes.

To explore the genomic context contributing most strongly to expression prediction, we prepared three distinct input datasets:

- 2.5 kb DNA sequence centered on the gene transcription start site (TSS);
- 2.5 kb DNA sequence upstream of the TSS;
- 2.5 kb DNA sequence downstream of the TSS.

We employed the Deep-Plant model pretrained on chromatin state data to predict the log1p (*log*(1 + *x*)) of gene expression levels from sequences. To adapt the model to RNA-seq–based gene expression prediction, the final output layer was modified to match the dimensionality of the RNA-seq expression profiles for each species. The remaining network weights were initialized from the pretrained model parameters, enabling transfer learning from chromatin-level regulatory features to transcript-level outcomes. Furthermore, we replaced the Softplus activation function in the output layer with ReLU, which yielded improved validation performance in regression tasks.

Depending on the experiment, pretrained weights were either fine-tuned or kept frozen to assess representational robustness. We trained the expression model using the standard Huber loss between predicted and observed expression levels. The model was optimized using AdamW, with a fixed learning rate *lr* = 5*e*^*−*5^. To prevent overfitting, we employed early stopping, terminating training if no improvement was observed for ten consecutive epochs on the validation dataset.

#### Attribution with In-silico-mutagenesis

We performed in silico saturation mutagenesis (ISM) on a test set of 3,843 *Arabidopsis thaliana* genes. For each gene, we generated all possible single-nucleotide variants (SNVs) within a 2.5 kb genomic window centered on the transcription start site (TSS). Each mutated sequence was evaluated using the fine-tuned gene expression model. For a mutation at position *i* to nucleotide *j* ∈ {*A, C, G, T*}, regulatory impact was quantified as the average cumulative *log*_2_ fold change (LFC) across all 3,142 predicted tissue- and condition-specific outputs:

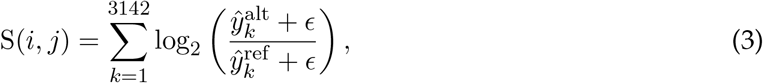

where 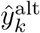 and 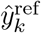 denote predicted expression for the alternative and reference sequences, respectively, for the *k*^*th*^ experiment. A constant *ϵ* = 0.5 was added to stabilize fold-change estimates at low expression levels.

Furthermore, ISM scores were normalized by subtracting, at each position, the mean across all four nucleotides. For each gene, mutations were ranked by their ISM score. High-impact variants were defined relative to gene-specific extremes. Up-regulatory mutations were those with positive cumulative LFC *S* ≥ 90% of the maximum positive LFC observed for that gene, whereas down-regulatory mutations had negative cumulative LFC with absolute magnitude *S* ≥90% of the maximum negative LFC.

#### A case study: the DREB1 gene family in Arabidopsis

We performed in silico saturation mutagenesis (ISM) on the three DREB1 genes separately across six experiments (see Supplementary Table 2 for more details). For each gene and experiment, ISM scores were computed and normalized at each position by subtracting the mean across nucleotides. High-impact positions were defined as those with scores ≥ 40% of the maximum observed score.

To identify TFs affected by these mutations, 20 bp sequences centered on each variant were scanned against JASPAR PWMs (2024 release). Motif scores were normalized to each PWM’s theoretical maximum. Mutations were classified as motif-disrupting if they reduced the motif score from ≥ 80% to ≤50% of its maximum.

To validate TF occupancy, we intersected mutation loci with available ChIP-seq BED files for the corresponding TFs. Motifs lacking supporting ChIP-seq peaks at the mutation site were excluded. If a variant disrupts a motif, the reference nucleotide was highlighted in red.

When visualizing *S*, the height at each position corresponds to the normalized ISM score of the reference nucleotide *j*_*ref*_ relative to the four possible nucleotides.

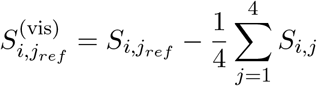

#### Fine-tuning Deep-Plant for enhancer activity in Arabidopsis

To further assess generalization across regulatory tasks, we fine-tuned Deep-Plant for enhancer activity prediction using published STARR-seq data in Arabidopsis [41]. The dataset comprised 38,901 training, 3,572 validation, and 3,442 test sequences, totaling 45,915 genomic regions (including 4,327 enhancers and 41,588 non-enhancer controls). Negative samples were selected to match the genomic distribution of enhancer loci: 53% exonic, 38% promoter-proximal (mostly ≤1kb from promoters), 6% in distal intergenic regions, and 1% within intronic, downstream ( ≤1kb from genes), or UTR regions (see Supplementary Figure 3 for more details). DNA sequences were extended to a fixed length of 2,500 bp. This was achieved by centering the window on each enhancer region and extending the coordinates upstream and downstream to reach the target length.

As in the gene expression task, we initialized training with pretrained Deep-Plant weights and fine-tuned (or froze) them as needed. For classification, the output activation function was changed from Softplus to Softmax, allowing probabilistic modeling of enhancer activity. We trained the enhancer model using the standard cross-entropy loss and AdamW optimizer with a fixed learning rate *lr* = 1*e*^*−*4^. Training was stopped if no improvement was observed for five consecutive epochs on the validation dataset.

To interpret the biological relevance of the Deep-Plant predictions, we first characterize the underlying complexity of the *Arabidopsis thaliana* enhancer datasdet landscape. Rather than a simple binary classification of active versus inactive, the dataset (published by Yongjun et al. [41]) stratifies enhancers into four distinct clusters. For data homogeneity and to facilitate interpretation, we only considered enhancers in Cluster 1 (C1, 37% of the total enhancers sequences), defined by high accessibility and enrichment for active histone acetylation marks (e.g., H3K9ac, H3K27ac) and H3K4 methylation, and physically associated with the most highly expressed genes in the genome. We kept a similar genomic distribution in negative samples and a 1:10 ratio of positive to negative samples. We evaluated a transfer learning approach based on logistic regression (LR) applied to the predicted epigenomic features generated by the pretrained chromatin-state model. In this setup, the entire pretrained Deep-Plant model was frozen and used as a feature extractor to obtain a chromatin state descriptor for each input sequence. Since the target ranges differ, chromatin predictions were mean-normalized prior to logistic regression to ensure all features have a similar scale. We also excluded the bias and used an L1 penalty to ensure relevant and interpretable weights. These feature representations were then used to train a downstream logistic regression classifier to predict enhancer activity. To assess the relative contribution of different regulatory modalities, we trained separate classifiers using different subsets of predicted targets.

1. **All targets:** utilized all 2,835 chromatin profiles predicted by Deep-Plant as input features.
2. **TF model:** included only 1,546 outputs corresponding to transcription factors and other DNA-binding proteins. We excluded experiment SRX3652551, which utilizes the human ten–eleven translocation methylcytosine dioxygenase 1 (TET1) for epimutagenesis in Arabidopsis, because TET1 is not a native plant protein.
3. **HM model:** comprised 1,119 outputs related to histone modifications.

We trained the Logistic regression models using Deep-Plant outputs, using the cross-entropy loss, L1 penalty (*λ* = 0.001), and AdamW optimizer with a fixed learning rate *lr* = 1*e*^*−*3^.

#### Fine-tuning Deep-Plant for detection of enhancer activity in corn

To assess Deep-Plant generalization across species, we also fine-tuned Deep-Plant for enhancer activity prediction using published STARR-seq data in the *Zea mays* inbred line B73 [71]). The pos-itive set comprises 1,718 enhancer regions experimentally identified by Oka et al [58]. To construct a robust negative set and prevent the model from learning trivial distinctions between coding and non-coding sequences (i.e., shortcut learning), we implemented a strict intergenic filtering strategy. Utilizing the standard B73 GFF3 annotation file, we explicitly masked all genomic regions corresponding to gene bodies (including Exons, Introns, and UTRs) and the enhancer regions. Negative samples were then randomly sampled exclusively from the remaining intergenic background. To reflect the realistic genomic scarcity of functional enhancers compared to the vast noncoding background, we imposed a 1:10 positive-to-negative ratio. Ultimately, the dataset comprised 16,082 training, 1,584 validation, and 1,232 test sequences, totaling 18,898 genomic regions (including 1,718 enhancers and 17,180 non-enhancer sequences). We also extended the sequences to a fixed length of 2,500 bp.

We utilized two variants of the DeepPlant model, pre-trained on well-annotated model organisms, to evaluate the impact of evolutionary distance on feature transferability:

- DeepPlant-AT: Pre-trained on the dicot model organism Arabidopsis thaliana.
- DeepPlant-OS: Pre-trained on the monocot model organism Oryza sativa (Rice).

Training was initialized with pretrained Deep-Plant weights, adapting the architecture by replacing the Softplus output activation with a Softmax layer. The model was fine-tuned using a standard cross-entropy loss objective and the AdamW optimizer with a fixed learning rate of 1*e*^*−*4^.To prevent overfitting, Training was stopped if no improvement was observed for five consecutive epochs on the validation dataset.

### 4.4 Benchmarking against DNA Language Models

#### AgroNT

AgroNT [14] is a DNA large language model tailored for plant genomics, trained on the DNA sequences of 48 plant species using a masked language modeling objective. AgroNT provides embeddings that can be fine-tuned for downstream predictive tasks such as regulatory element classification or gene expression modeling.

#### PDLLM

Plant DNA Large Language Model PDLLM is a suite of lightweight, DNA language models specifically tailored for plant and crop genomics [15]. They were explicitly developed to solve the computational accessibility problem posed by massive models like AgroNT, offering an alternative that can be fine-tuned on commodity GPUs. We used their Plant-DNAGPT-6mer model.

#### Training of AgroNT and PDLLM

Model training and evaluation utilized the exact datasets and sample splits as the DeepPlant setup. All models were fine-tuned for 15 epochs using the IA^3^ technique [72], with optimal models selected based on minimum validation loss. While full training was conducted on an NVIDIA A100 GPU (80 GB), training times were benchmarked on an NVIDIA RTX 4090 (24 GB) for fair comparison. Task-specific heads and loss functions were assigned as follows: a regression head with Poisson loss for chromatin state, a regression head with Mean Squared Error (MSE) loss for gene expression, and a classification head with crossentropy loss for enhancer activity.

## Acknowledgments

This material is based upon work supported by the U.S. Department of Energy, Office of Science, Biological and Environmental Research program (BER), under Award Number DE-SC-0024459.

## Data availability

The code repository for training RNA-seq deep learning models, including example code to use the model as well as scripts for variant scoring, is available at https://github.com/Addaoud/DeepPlant. The repository also contains code relevant to the analyses and results presented in the manuscript. Pre-trained Deep-Plant model weights are available through HuggingFace.

