## Supplemental text and figures for "Deep-Plant: a supervised foundation model for plant regulatory genomics"

| Experiment type | Specific factor type | Arabidopsis | Rice |
|---|---|---|---|
| DNA-binding proteins | Transcription factor | 1,243 | 49 |
| | DNA-binding protein | 202 | 0 |
| | RNA polymerase | 76 | 24 |
| | RNA polymerase subunit | 25 | 0 |
| | Methylase | 0 | 1 |
| | Cryptochrome | 1 | 0 |
| Histone and Histone-related | Histone methylation | 643 | 186 |
| | Histone acetylation | 196 | 57 |
| | Histone-related | 265 | 28 |
| | histone ubiquitination | 7 | 0 |
| | Heterochromatin | 8 | 0 |
| DNase and ATAC-seq | – | 164 | 5 |
| DNA-methylation | – | 5 | 0 |

Table 1: Detailed epigenomic data in Arabidopsis and rice. We collected all available datasets from ChIP-Hub and averaged replicates.

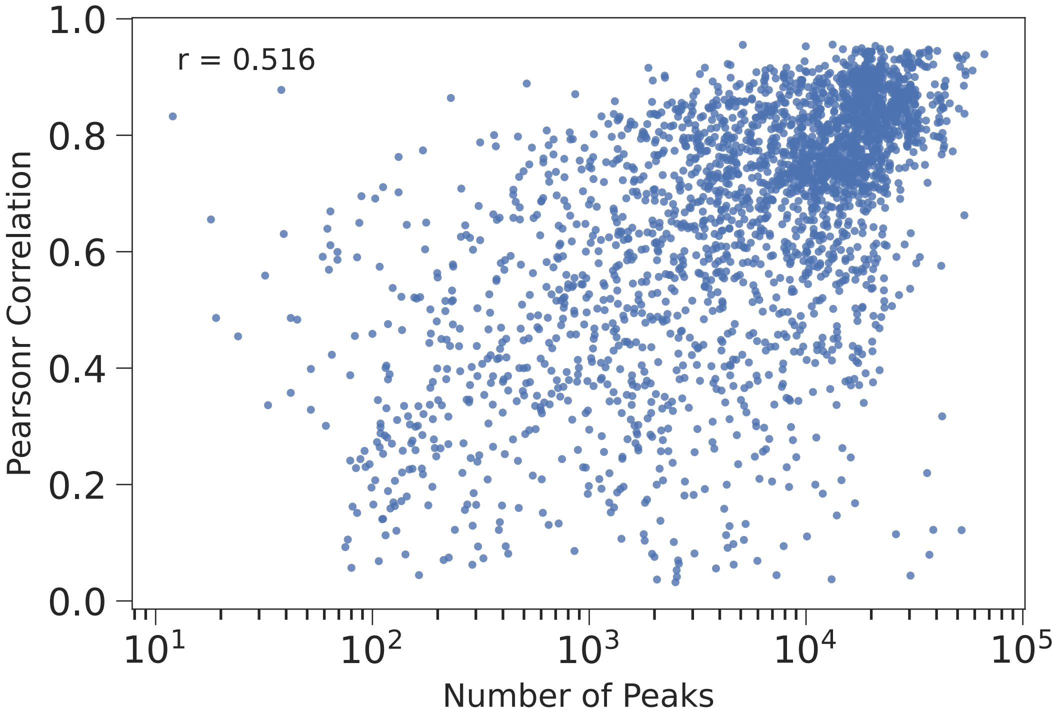


Figure 1: Correlation between model performance and peak density. Scatter plot showing the relationship between the number of peaks identified in the experimental BED files (x-axis, $\log_{10}$ scale) and the Pearson correlation coefficient achieved by DEEP-PLANT (y-axis).

### Biological Interpretation of Convolutional Neural Network Filters

To elucidate the biological function of sequence features learned by our model, we established a systematic computational pipeline designed to link each filter in the model's first convolutional layer to known transcription factor (TF) binding motifs. This pipeline consists of three main stages: first, identifying the DNA sequences that maximally activate each filter; second, performing de novo motif discovery on these sequences ; and third, comparing the discovered motifs against a database of known Arabidopsis TF binding motifs for annotation.

### Identification and Extraction of Maximally Activating DNA Sequences

Our analysis focuses on the 512 filters of DeepPlant's first convolutional layer. The input data consisted of 2,500 bp genomic sequences from the training set, centered around transcription start sites (TSS). These sequences were transformed into four-channel one-hot encoded matrices, a standard representation for deep learning models processing nucleotide data. For each filter, we computed its activation score across the entire training dataset by passing all sequences through the model and recording the maximum activation value produced by the filter for each sequence. We then identified the global maximum activation value for that filter across all sequences and set a screening threshold at 80% of this value. Any sequence eliciting a maximum activation that met or exceeded this threshold was classified as a high-activating sequence. Subsequently, from each high-activating sequence, we precisely located the position of maximum activation and extracted the DNA subsequence corresponding in length to the filter's kernel size. To construct a representative set of motifs, these extracted subsequences were deduplicated; if the same sequence was extracted multiple times, only the instance associated with the highest activation score was retained. This process yielded a dedicated FASTA file for each of the 512 filters, containing its corresponding set of high-activating, non-redundant DNA sequences.

### De Novo Motif Discovery

The sets of high-activating sequences for each filter were subsequently used for de novo motif discovery, a task for which we employed the STREME algorithm (Bailey, 2021). STREME is designed to identify statistically significant, ungapped sequence patterns (motifs) that are enriched within a given set of sequences. For the STREME analysis, motifs were constrained to a length of 8 to 15 nucleotides, a reporting p-value threshold of 0.05 was used, and motifs were centrally aligned. This procedure resulted in the discovery of a set of potential regulatory motifs for each convolutional filter, which were stored in MEME format.

### Motif Annotation and Transcription Factor Matching

To functionally annotate the discovered motifs, we utilized the TOMTOM algorithm (Gupta et al., 2007) to perform a quantitative comparison against a database of known TF binding motifs. In this step, we extracted only the most statistically significant (i.e., the top-ranked) motif from the STREME output for each filter to serve as the query. This query motif was then compared against the ArabidopsisDAPv1 reference database, which contains a comprehensive collection of experimentally validated TF binding profiles for Arabidopsis thaliana derived from DNA affinity purification sequencing (O'Malley et al., 2016). The key parameters for the TOMTOM comparison were configured to use Pearson correlation as the similarity metric, require a minimum overlap of 5 positions between the query and target motifs, and set an E-value threshold of 10.0 for reporting matches. The final output of this pipeline is a ranked list of known TF binding motifs that are statistically significant matches for the motif learned by each convolutional filter, providing a direct annotation and functional hypothesis for the biologically relevant patterns learned by the neural network from DNA sequences.

#### Convolutional Filters Learn Biologically Relevant Features Matching Known Transcription Factor Motifs

To interpret the biological function of the features learned by the deep learning model, we performed a systematic functional annotation of all 512 filters in the first convolutional layer of the arabidopsis Deep-

PLANT model. By matching the sequence patterns activated by each filter against a comprehensive database of Arabidopsis transcription factor (TF) binding motifs, we found that 500 filters (97.65%) successfully matched one or more known TF binding motifs with adjusted p-values below 0.05 (see Fig. 2). This result strongly demonstrates that the model is not an uninterpretable "black box," but rather is capable of autonomously learning and identifying authentic biological features that are directly related to gene regulation. We assessed the statistical significance of all filter-motif matches. The distribution of -log10(p-values) is heavily skewed toward larger values, with the vast majority of matches having a p-value far below 0.01. This confirms that the similarity between the model's learned sequence patterns and the TF binding motifs in the database is highly reliable and statistically significant.

To further characterize the regulatory site learned by the model, we performed a categorical analysis of the TF families represented by the 512 matched motifs. The results show that the learned features cover a wide variety of TF families. Among them, the three most frequently identified families were MYB (matched by 81 filters), MYBrelated (48 filters), and C2C2dof (42 filters). These families are known to play critical roles in numerous core biological pathways in Arabidopsis, including growth, development, stress response, and hormone signaling, indicating that our model successfully captured these essential regulatory elements.

To visually illustrate the degree of similarity between the learned and known motifs, we selected several representative examples for comparison (Figure 2). These examples were chosen to highlight both statistical significance and family representation. Overall, the learned motifs discovered by the model show concordance with the sequence logos of their best matches from the database. For instance, Filter 21 provided one of the most statistically significant matches (q-value $= 1.7 * 10^{-5}$) and accurately captured the binding site for the MADS family of transcription factors (Figure 2). This plant-specific family play important roles in flower development from the early step of determining the identity of floral meristems [1]. Two other highly significant matches also showcase the breadth of the model's learned features. Filter 14 (q-value $= 2.11 * 10^{-4}$) matched the G2-like family, a subfamily of the GARP superfamily, associated with chloroplast maturation influence ozone resilience via the modulation of stomatal behavior[2]. Similarly, Filter 392 (q-value $= 3.88 * 10^{-4}$) represents the MYB-related family, which plays a key role in regulatory networks controlling development, metabolism and responses to biotic and abiotic stresses [3]. Together, these examples demonstrate that the model not only learned statistically robust features but that these features also correspond directly to key biological functions in plant growth, metabolism, and development. In summary, our findings clearly show that the filters of DEEP-PLANT's convolutional layer can serve as highly effective detectors of biological features. The model autonomously learned a large vocabulary of authentic DNA sequence patterns relevant to transcriptional regulation, and these patterns cover a diverse range of critical transcription factor families, all validated by rigorous statistical testing. This not only confirms the biological relevance of our model but also establishes a powerful framework for using deep learning to decipher the sequence-level code of gene expression.

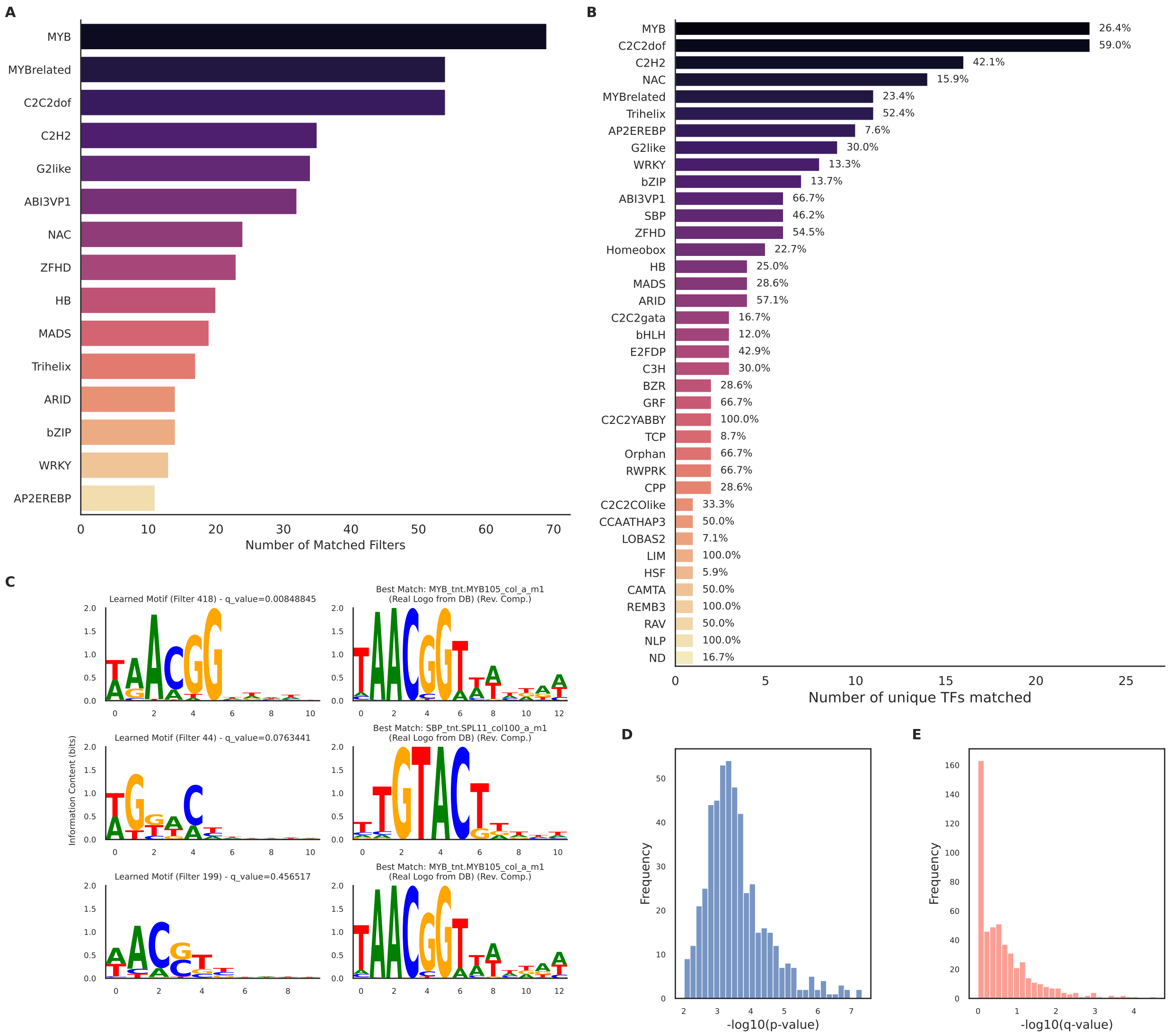


Figure 2: Filter analysis Mapping of DEEP-PLANT convolutional filters to known transcription factor (TF) families in Arabidopsis. Bar plot showing the number of DEEP-PLANT filters whose learned sequence motifs significantly match known transcription factor binding motifs from Arabidopsis. Each bar represents a TF family (number of TFs within that family is indicated), illustrating the diversity of regulatory patterns captured by the model and highlighting families such as WRKY, bZIP, and MYB as among the most frequently recovered. We also show the examples of mapped filters to known Arabidopsis motifs, and the distribution of p-values and q-values for the significance of the mapped filters.

| Experiment accession | Condition | Tissue |
|---|---|---|
| ERX1268170 | Normal | Leaves |
| SRX1881751 | Cold 1 hour | Leaves |
| SRX1881753 | Cold 3 hour | Leaves |
| SRX1881761 | Cold 6 hour | Leaves |
| SRX1881839 | Cold 12 hour | Leaves |
| SRX1881841 | Cold 24 hour | Leaves |

Table 2: Gene expression experiments used for in-silico-mutagenesis analysis of the DREB1 genes in *Arabidopsis thaliana*.

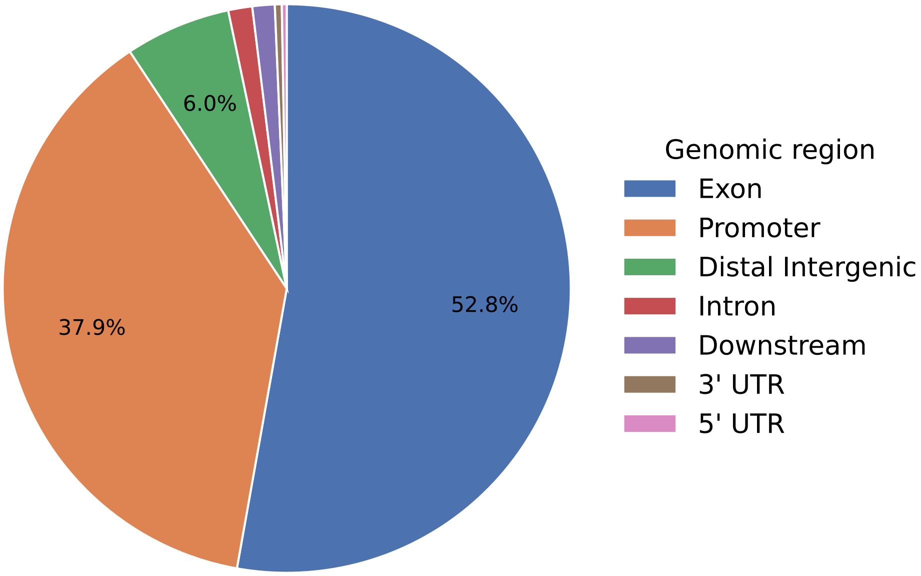


Figure 3: Genomic distribution of enhancers. Pie chart showing the proportion of enhancers overlapping exons, promoters, distal intergenic regions, introns, downstream regions, and untranslated regions (UTRs). Percentages indicate the fraction of enhancers in each genomic category.
